# Postural test to differentiate primary aldosteronism from low-renin hypertension

**DOI:** 10.1101/2025.05.16.654625

**Authors:** Irene Tizianel, Elena Pagin, Eugenio Ragazzi, Alberto Madinelli, Simona Censi, Chiara Sabbadin, Franco Mantero, Caterina Mian, Mattia Barbot, Giorgia Antonelli, Filippo Ceccato

## Abstract

**Background:** The diagnostic accuracy of screening and confirmatory tests to differentiate primary aldosteronism (PA) among patients with low-renin hypertension (HTN) is suboptimal. We aimed to assess the role of postural stimulation test (PST, previously used for PA subtyping) in differentiating PA from low-renin HTN.

**Patients and methods:** Clinical and endocrine data in clinostatic position (CP) and orthostatic position (OP) during PST were evaluated in 190 hypertensive patients: 80 PA and 110 low-renin HTN. Multivariate techniques were computed: Principal Component Analysis (PCA), Partial Least Square-Discriminant Analysis (PLS-DA) and k-means clustering.

**Results:** PST response differentiated our cohort: 96% of PA were detected in the 56/190 patients with always suppressed renin levels, 80% of patients with low-renin HTN were identified among 56/190 subjects with de-suppression of renin from CP to OP and 78/190 with always measurable renin. Increased potassium and measurable renin in OP were predictors of low-renin HTN. Cluster analysis distinguished PA from low-renin HTN: Cluster 2 included 104/110 low-renin HTN; Cluster 1 PA patients showed a higher frequency of suppressed renin levels at baseline and during PST (100% in CP and 95% in OP, respectively). Cluster 1 low-renin HTN patients had lower potassium and a higher frequency of suppressed renin levels at diagnosis and during PST, compared to Cluster 2. PLS-DA and PCA confirmed that renin in OP, renin response to PST and presence of hypokalemia were the most relevant parameters for distinguishing PA from low-renin HTN.

**Conclusion:** Renin response during PST can be used to differentiate PA from low-renin HTN.

## Introduction

Primary aldosteronism (PA) is the most common cause of secondary hypertension (HTN): it is characterized by autonomous aldosterone secretion, independent from renin, sodium levels, and volemic status [1]. Unilateral adrenalectomy or medical treatment with mineralocorticoid receptor antagonists significantly ameliorate PA outcomes: a correct approach upon reliable diagnostic tests would be of great value [2]. PA diagnosis requires different steps: initial screening is based on measurement of aldosterone to renin ratio (ARR). An increased ARR enhances the likelihood of PA, however it can reflect a low renin status, thus failing to recognize an underlying autonomous aldosterone secretion [3]. Therefore, additional tests should be performed to confirm or exclude PA. Among them, the saline infusion test (SIT, which relies on aldosterone suppression in response to volume expansion) and the captopril challenge test (CCT, based on aldosterone reduction after angiotensin converting enzyme inhibitor), are routinely used worldwide [4], [5]. Recognizing PA is crucial: excessive mineralocorticoid receptor activation due to aldosterone excess is associated with higher cardiovascular morbidity and mortality compared to essential HTN. Therefore, an appropriate and timely recognition and treatment can mitigate this risk [6], [7], [8]. Traditionally, PA is defined as a unilateral or bilateral form according to the subtype (unilateral adenoma or bilateral adrenal hyperplasia). However, it is no longer considered as a dichotomous condition, but rather a continuum of disease featured by renin-independent aldosterone excess [6], [9], [10]. Low-renin HTN is a common condition which counts for at least 20-30% of hypertensive patients [11]. Low renin levels, arbitrarily defined as direct renin concentration (DRC) from 4 to 12 mIU/L or plasma renin activity (PRA) <0.5 to 1 ng/mL·h are detectable in more or less one quarter of hypertensive patients, with higher prevalence in the elderly and in subjects of African origin [3], [12]. The postural stimulation test (PST) was used in the past decades to distinguish different subtypes of PA (unilateral from bilateral forms). However, results were contradictory, and some studies reported the inability of PST to discriminate between the two different subtypes [13], [14]. Recently, PST was proposed as a diagnostic test for PA confirmation, particularly in patients with a high likelihood of final PA diagnosis [15]. The principal aim of our study is to assess the role of PST in differentiating PA from patients with low-renin HTN.

## Material and methods

### Study design

We conducted a single-centre observational retrospective study at Padova University-Hospital, approved by local Ethics Committee: PITACORA, Pituitary Thyroid Adrenal Tumors Cardiovascular and Outcome related long-term Assessment (protocol number AOP3318, registration 5938-AO-24). Data are available in the Repository of the University of Padova [16]. This observational study was conducted following the STrengthening the Reporting of OBservational studies in Epidemiology (STROBE) guidelines [17].

### Patient selection

We retrospectively analyzed a cohort of 226 patients followed at Padova University-Hospital from January 2009 to July 2024. These subjects underwent endocrinological evaluation for arterial HTN associated with at least one positive aldosterone/renin ratio (ARR) performed with a proper washout from interfering medications, and concomitant aldosterone levels >10 ng/dL (277 pmol/L). To define pathological ARR we adopted a cut-off value of 30 with PRA (ng/mL/h) and 91 with DRC (mU/L), according to 2016 Endocrine Society Clinical Practice Guideline [1]. Thirty-six patients were excluded for incomplete data: the final cohort consisted in 190 patients who underwent at least one confirmatory test (SIT/CCT or both). An aldosterone post-SIT (performed in seated position) >10 ng/dl (277 pmol/L) and/or an aldosterone suppression <30% after CCT confirmed PA diagnosis. We adopted 10 ng/L as the aldosterone post-SIT threshold to reduce false negative results, since SIT is a confirmatory test and therefore a higher specificity is required. In the “grey zone” indicated by aldosterone levels post-SIT between 5 and 10 ng/L (138-277 nmol/L) we classified the patient according to CCT results: PA was confirmed in case of positive CCT. Considering the not negligible rate of false negative results of SIT, we preferred to combine CCT as a second confirmatory test. Only 9 patients underwent SIT alone, however, they all had aldosterone post-SIT > 10 ng/dl (277 pmol/L) and therefore were classified as PA (see Figure 1). In case of positive confirmatory tests patients were diagnosed with PA (n=80), and PA subtyping was pursued in 55 cases: 34 were classified as unilateral and 21 with bilateral disease. Adrenal venous sampling (AVS) was not performed in 25 subjects due to patient’s refusal or poor surgical eligibility. Between patients with unilateral PA, according to Primary Aldosteronism Surgery Outcome (PASO) criteria [18], 44% had complete clinical and biochemical success, while 56% had partial clinical and complete biochemical success. In case of negative confirmatory tests, the patient was classified as low-renin HTN. Baseline clinical and biochemical data at diagnosis were assessed. PST consisted in aldosterone and renin measurement in clinostatic position (CP, after resting in supine position for at least an hour and a half) and subsequently in orthostatic position (OP, after 2 hours of walking). Since the limited reliability of renin measurements in the lower part of the reference range and the unreliable and non-unique conversion factor from PRA to DRC [19], we decided to consider renin levels not as a continuous variable but as a categorical one: measurable vs suppressed, thus above or below the lower limit of normality, respectively. Suppressed renin was defined as PRA <1 ng/mL·h or DRC <2 mIU/L. All biochemical analyses were measured in the ISO15189 accredited Laboratory of the University-Hospital of Padova. From 2009 to 2014, PRA and serum aldosterone were measured using radioimmunoassay kit (RIA, Beckman Coulter kit). From 2015, DRC and serum aldosterone were measured using chemiluminescence immunoassays on a completely automated system (DiaSorin, LIAISON® XL instrument) with LIAISON® Direct Renin kit (DiaSorin, Saluggia, Italy) and LIAISON® XL Aldosterone kit, respectively. The concordance between RIA and chemiluminescence immunoassays was verified, as previously reported [20].

**Figure 1.**
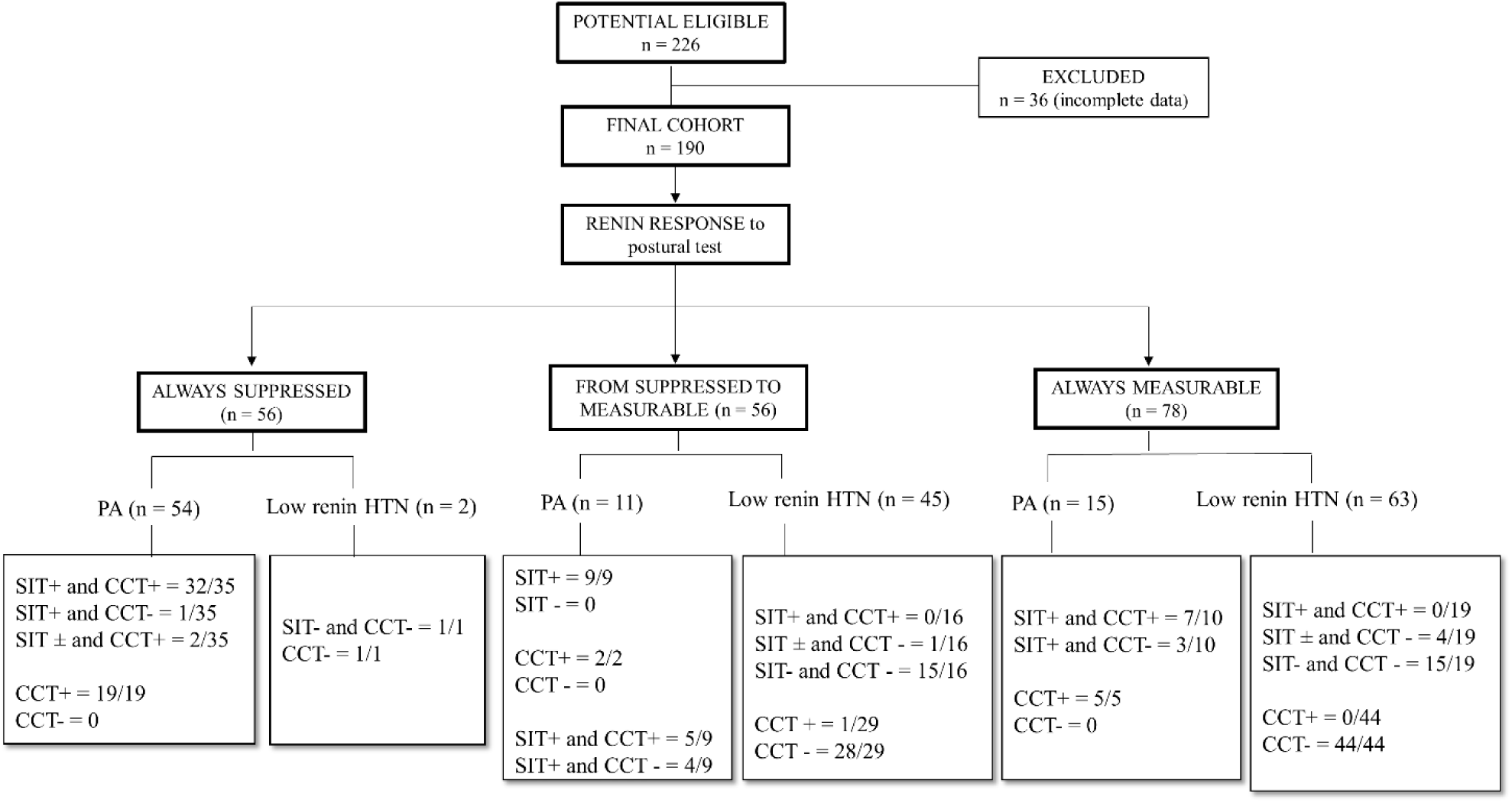
Diagnostic flowchart according to renin response during postural test. Abbreviations: CCT captopril challenge test, SIT saline infusion test, PA primary aldosteronism, HTN hypertension, Response to SIT: +: positive (aldosterone after SIT >10 439 ng/dL, 277 pmol/L); ± aldosterone after SIT in the grey zone 5-10 ng/dL (138-277 pmol/L); -: SIT negative (aldosterone after SIT <5 ng/dL, 138 pmol/L); CCT: + reduction of aldosterone post CCT <30%; -: reduction of aldosterone after CCT >30%.

### Statistical analysis

Categorical variables were reported as frequencies and percentages, while continuous ones were reported as median and interquartile range (IQR). Nonparametric tests were used to compare groups. Data analysis was performed using SPSS version 24 software package for Windows (SPSS Inc), JMP Version Pro 17 software for Windows (SAS Institute Inc), and MetaboAnalyst 6.0 [21]. Statistical significance was accepted at p<0.05. The Fisher exact test, with Bonferroni correction for multiple comparisons, was used to compare proportions between groups. Mann Whitney U-test, or Kruskal-Wallis test, were used for comparison of continuous variables, between two or three independent groups, respectively. Univariate and multivariate logistic regression were conducted to assess the association between the final diagnosis as dependent variable, and clinical and biochemical variables as predictors. Adjusted logistic regression models were based on clinical relevance, collinearity and interactions. Nonparametric multivariate methods were used to obtain an integrated analysis. Categorical data were pre-processed as dummy code. Since zero values or missing values cause difficulties in the analysis, the software replaced zero values with a conventional value; missing values were estimated using the K-Nearest Neighbours method. Volcano plot (significance vs fold-change) was obtained to illustrate the variables’ relevance. Among nonparametric multivariate techniques, principal component analysis (PCA) unsupervised method was used, permitting reduction of the dimensions of the data set and highlighting relevant variables. By combining the original data, PCA provides a summary based on fewer derived variables (the scores), whose profiles are represented by the loadings, which indicate the contribution to the principal components [22]. k-means clustering was used to ascertain patients’ subdivision based on their similarity according to minimum distance from the centroid of k clusters, by using the available variables [23]. The supervised classification method of partial least squares discriminant analysis (PLS-DA) was also used to reduce data dimensionality and provide further information about the profile of data, such as the identification of most relevant features [24]. Additionally, hierarchical cluster analysis was performed with Euclidean distance measure and using Ward clustering algorithm.

## Results

### Clinical and biochemical data at diagnosis

Baseline clinical and endocrine data of the two groups are reported in Table 1. Age distribution was similar, while female patients were prevalent in the low-renin HTN group. As expected, aldosterone levels at baseline were higher in PA compared to low-renin HTN (855 vs 484 pmol/L, p<0.001) and PA patients had lower potassium levels (3.1 vs 3.9 mmol/L, p<0.001). Also, prevalence of renin suppression at baseline was higher in PA compared to low-renin HTN (59 vs 25%, p<0.001). BMI and glucose metabolism alterations (GMAs) distribution were similar in the two groups. PA patients showed resistant HTN more frequently (28% vs 15%, p=0.043).

**Table 1.** Baseline clinical, biochemical and postural test data in the two groups. Data are presented as median and IQR (in brackets) or frequencies and percentages (in brackets). Abbreviations: BMI: body mass 533 index, HTN: hypertension, GMA: glucose metabolism lterations; CP: clinostatic position; OP: orthostatic position; ARR: aldosterone to renin ratio. Hypokalemia: K+ levels < 3.5 mmol/L. a) Comparison between the 3 subgroups of PA; b) Comparison between the 3 subgroups of low-renin HTN 412

|  |  | PA (n = 80) |  | low-renin HTN (n = 110) | p value |
| --- | --- | --- | --- | --- | --- |
|  |  | At diagnosis |  |  |  |
| Age (years) |  | 50.5 (43.7-58.5) |  | 53 (42.2-5-62.5) | 0.217 |
| sex (% female) |  | 37/80 (46%) |  | 69/110 (63%) | 0.027 |
| Aldosterone (pmol/L) |  | 855 (604-1223.7) |  | 484 (395.2-627.5) | <0.001 |
| Renin (suppressed/tot) |  | 47/80 (59%) |  | 27/110 (25%) | <0.001 |
| ARR |  | 323.5 (219-426) |  | 149.3 (107.2-205.4) | <0.001 |
| K<sup>+</sup> levels (mmol/L) |  | 3.1 (2.8-3.6) |  | 3.9 (3.5-4.1) | <0.001 |
| hypokalemia (yes/tot) |  | 55/79 (69%) |  | 16/97 (16%) | <0.001 |
| BMI (kg/m<sup>2</sup>) |  | 26 (23-28) |  | 27 (24-30) | 0.055 |
| HTN drugs (> 3 drugs) |  | 22/80 (28%) |  | 16/110 (15%) | 0.043 |
| GMA (normal/tot) |  | 53/73 (73%) |  | 73/97 (75%) | 0.925 |
|  |  | During postural test |  |  |  |
| Renin – CP (suppressed/tot) |  | 65/80 (81%) |  | 47/110 (43%) | <0.001 |
| Renin – OP (suppressed/tot) |  | 55/80 (69%) |  | 2/110 (2%) | <0.001 |
| Renin from suppressed to measurable (n/tot) |  | 10/80 (12%) |  | 45/110 (40%) | <0.001 |
| Aldosterone – CP (pmol/L) |  | 545 (419-888) |  | 249 (180-368.5) | <0.001 |
| Aldosterone – OP (pmol/L) |  | 946 (633.75-1361.5) |  | 600 (432-878.75) | <0.001 |
| Cortisol – CP (nmol/L) |  | 258.5 (195.5-324.25) |  | 257.5 (200.5-315) | 0.738 |
| Cortisol – OP (nmol/L) |  | 235 (179.75-320.75) |  | 258.5 (192-359) | 0.163 |
| Aldosterone increase (%) | entire cohort | 50.4 (6.7-103.7) |  | 126.9 (70-197) | <0.001 |
|  | renin (group always suppressed) | 38.3 (5.6-102.9) |  | 133 (81.9-389) | p = 0.680<sup>a</sup><br>p = 0.740<sup>b</sup> |
|  | Renin (group from suppressed to measurable) | 40.4 (-1.1-168) |  | 110.9 (74.8-186.6) |  |
|  | Renin (group always measurable) | 82.5 (20.1-103.7) |  | 132.8 (64.3-213.2) |  |

Abbreviations: BMI: body mass index, HTN: hypertension, GMA: glucose metabolism alterations; CP: clinostatic position; OP: orthostatic position; ARR: aldosterone to renin ratio. Hypokalemia: K<sup>+</sup> levels < 3.5 mmol/L. a) Comparison between the 3 subgroups of PA; b) Comparison between the 3 subgroups of low-renin HTN 412

### Postural stimulation test (PST)

The frequency of suppressed renin was higher in PA compared to low-renin HTN (in CP 81 vs 43%, p<0.001, in OP 69 vs 2%, p<0.001), as summarized in Table 1. Considering renin response to PST we focused on patients whose renin levels changed from suppressed in CP to measurable in OP: this category was more prevalent in patients with low-renin HTN compared to PA (40 vs 12%, p<0.001). Aldosterone levels (both in CP and in OP) were higher in PA, and the rate of aldosterone increase during PST was higher in low-renin HTN (50% in PA vs 127% in low-renin HTN, p<0.001), suggesting a preserved physiological axis. Choosing an arbitrarily threshold of aldosterone increase ≥50%, 40/80 PA and 97/110 low-renin HTN showed this behavior (50 vs 88%, p<0.001). Considering the PA cohort, the rate of aldosterone increase did not differ according to renin response from CP to OP. In low-renin HTN the percentage of aldosterone increase (≥50% as our selected criteria) is higher than PA because in the former the aldosterone still increases after renin stimulus from CP to OP, while in the latter the aldosterone levels are always high and independent from renin levels. Cortisol levels during PST were similar in the two groups and did not increase from CP to OP, supporting the idea that aldosterone response in PA is autonomous from renin and ACTH levels. Figure 1 shows the diagnostic flowchart according to renin response from CP to OP during PST. Overall 56/190 patients (29%) presented suppressed renin levels both in CP and in OP, 56/190 (29%) showed de-suppression of renin levels from CP to OP, and 78/190 (42%) had non-suppressed renin in CP which remained measurable in OP. In the arm of always suppressed renin in CP and in OP 54/56 (96%) patients were PA, while 45/56 (80%) of patients in the category from suppressed (in CP) to measurable (in OP) renin had low renin HTN as final diagnosis. Finally, 63 out of 78 (81%) of patients with measurable renin in CP and in OP were classified as low-renin HTN. Regarding patient’s distribution according to renin response to PST, the one patient with positive CCT in the renin response category from suppressed to measurable had an aldosterone decrease after CCT of 28% and a normal ARR after CCT, therefore he was considered as low-renin HTN. Instead, in the category of always measurable renin, the four patients with aldosterone after SIT in the “grey zone” and negative CCT had aldosterone levels after SIT of 6-7 ng/dl, and an ARR performed over the following 6-12 months was negative, therefore PA was excluded. In the category of always suppressed renin the two PA patients with negative SIT had an aldosterone after SIT in the “grey zone” (5.5 and 5.8 ng/L, respectively), but CCT was positive and therefore PA was confirmed. Univariate logistic regression analysis showed that the odds ratio (OR) of low-renin HTN as the final diagnosis was 4.37-fold higher in patients with measurable renin levels (95% CI 2.35-8.15); moreover, the OR for increasing potassium levels was 11.78-fold higher for ruling out PA (Table 2). In the multivariable regression analysis, aldosterone levels at diagnosis, serum potassium levels, and renin in OP during PST remained significant predictors of the final diagnosis (Chi-Square=165.94, df=5 and p<0.001). All five predictors “explain” 82% (Nagelkerke R2) of the variability of the final diagnosis of PA. The model correctly predicted 86% of cases of PA and 97% of cases of low-renin HTN, giving an overall percentage correct prediction rate of 92%. In particular, multivariable regression analysis confirmed that the odds for low-renin HTN as final diagnosis were 12-fold higher for increasing potassium levels and 352-fold higher for measurable renin in OP during PST. Increasing aldosterone levels at diagnosis, as shown in univariate analysis, were less informative in predicting the final diagnosis, thus showing a OR 1-fold lower for the diagnosis of low-renin HTN (summarized in Table 3).

**Table 2.** Univariable binary logistic regression analysis for predicting low-renin HTN as final diagnosis.

|  | OR | 95% CI | *p* value |
| --- | --- | --- | --- |
| **At diagnosis** |  |  |  |
| **Aldosterone (pmol/L)** | 0.988 | 0.987-0.999 | < 0.001 |
| **Measurable renin** | 4.37 | 2.35-8.15 | < 0.001 |
| **ARR** | 0.988 | 0.985-0.982 | < 0.001 |
| **K<sup>+</sup> levels (mmol/L)** | 11.78 | 5.56-24.94 | < 0.001 |
| **During postural test** |  |  |  |
| **Measurable renin in CP** | 5.81 | 2.95-11.42 | < 0.001 |
| **Measurable renin in OP** | 116.64 | 26.63-510.58 | < 0.001 |
| **Renin always measurable** | 0.93 | 0.38-2.26 | 0.879 |
| **Renin always suppressed** | 0.008 | 0.002-0.04 | < 0.001 |
| **Aldosterone increase (%)** | 1.00 | 0.99-1.00 | 0.124 |

**Table 3.** Multivariable binary logistic regression analysis for predicting low-renin HTN as final diagnosis.

|  | OR | 95% CI | *p* value |
| --- | --- | --- | --- |
| At diagnosis |  |  |  |
| Aldosterone (pmol/L) | 0.997 | 0.995-0.998 | < 0.001 |
| Renin (mIU/l) | 2.26 | 0.55-9.17 | 0.254 |
| K<sup>+</sup> levels (mmol/L) | 12.47 | 3.36-42.5 | < 0.001 |
| During postural test |  |  |  |
| Measurable renin in CP | 1.15 | 0.30-4.36 | 0.830 |
| Measurable renin in OP | 351.64 | 31.35-3921 | < 0.001 |

### The big-data approach

A multivariate analysis was tested to assess the relationship between the considered variables. In Fig. 2A we depict the PCA loadings plot of categorized data: the direction of vectors representative of renin in CP, renin in OP, and renin response during PST indicate high mutual correlation. On the contrary, the relationship between metabolic parameters was limited: the angle between vectors of GMA, dyslipidemia, and BMI distribution was close to or greater than 90°, indicating a poor or negative correlation. Also, arterial HTN vector showed an angle close to 90°, confirming no correlation, as expected per selection criteria: all patients of the entire cohort had HTN. By means of an unsupervised approach, the non-hierarchical k-means clustering technique, it was possible to observe how PA and low-renin HTN subjects can be sorted in two clusters (Figure 3). The algorithm calculates each cluster’s mean, and if a datum is closer to the nearest mean, this datum becomes part of that cluster. The model was able to distinguish patients with low-renin HTN: Cluster 1 correctly included 57/80 patients with PA (63%) and Cluster 2 correctly included 104/110 patients with low-renin HTN (95%). Therefore, the model can identify almost all patients with low-renin HTN, who are homogeneous for the studied characteristics. We further analyzed the clinical and biochemical picture of PA and low-renin HTN patients as resulted according to Cluster 1 and Cluster 2 (Table 4.). Cluster 1 patients with PA showed an endocrine profile characterized by a higher frequency of suppressed renin levels at baseline and during PST, compared to Cluster 2, with a prevalence of suppressed renin in CP or OP of 100% and 95%, respectively. Furthermore, Cluster 1 low-renin HTN patients had lower potassium levels at diagnosis and a higher frequency of suppressed renin levels at baseline and during PST, compared to Cluster 2. Therefore an “hypokaliemic and renin suppressed phenotype” identified low-renin HTN patients of Cluster 1, suggesting an intermediate phenotype between patients with low-renin HTN and PA in Cluster 1. To further investigate the cohort, we conducted an additional unsupervised approach through the non-hierarchical k-means clustering technique, dividing the whole cohort into three clusters (Figure 4). Interestingly, this cluster analysis showed subgroups similar to those obtained with a priori classification by using the criteria “renin response during PST” depicted in Figure 1. In particular, Cluster 1 was made of 54 PA and 2 low-renin HTN and corresponded to the category of always suppressed renin, Cluster 2 consisted of 11 PA and 45 low-renin HTN, such as the category of from suppressed to measurable renin and Cluster 3 was made of 15 PA and 63 low-renin HTN like the category of always measurable renin. As depicted in Figure 2 Panel B, the Partial Least Square-Discriminant Analysis (PLS-DA) displayed that renin in OP, renin response to PST and presence of hypokalemia were the best parameters for distinguishing PA from low-renin HTN. If pathological, they showed high accuracy for PA diagnosis, conversely if normal they show low diagnostic accuracy for the diagnosis of low-renin HTN. The same findings were demonstrated by volcano plot (Figure 2 Panel C) in which renin in OP, renin response and hypokalemia emerged as the most significant predictors of the final diagnosis. Hierarchical clustering depicted in Figure 5 confirmed the same variables as the most indicative of PA if pathological, while in case of normal values PA could not be excluded in favor of low-renin HTN.

**Figure 2.**
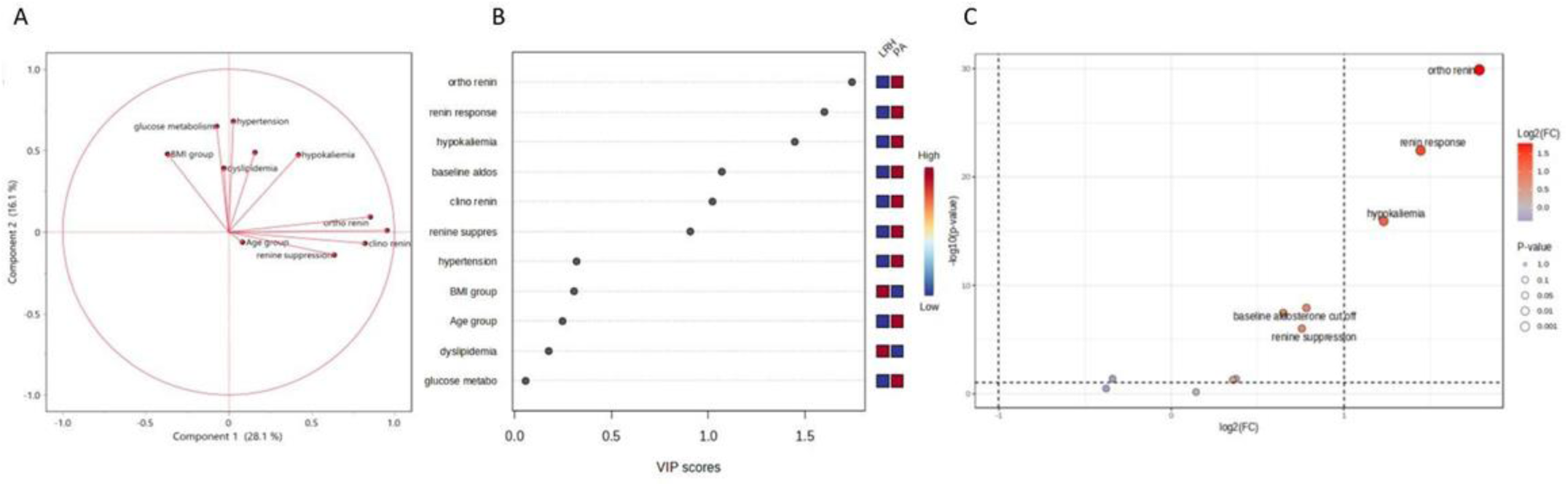
A) Loadings plot from PCA indicating the contribution of each parameter. B) Important features identified by Partial Least Square-Discriminant Analysis according to VIP scores. The colored boxes on the right indicate the relative value of the corresponding parameter in each group under study. C) Volcano plot. Important features selected by volcano plot with fold change threshold (x axis) 2 and t-test threshold (y axis) 0.1. The red circles represent features above the threshold. Both fold chances and p values are log transformed.

**Figure 3.**
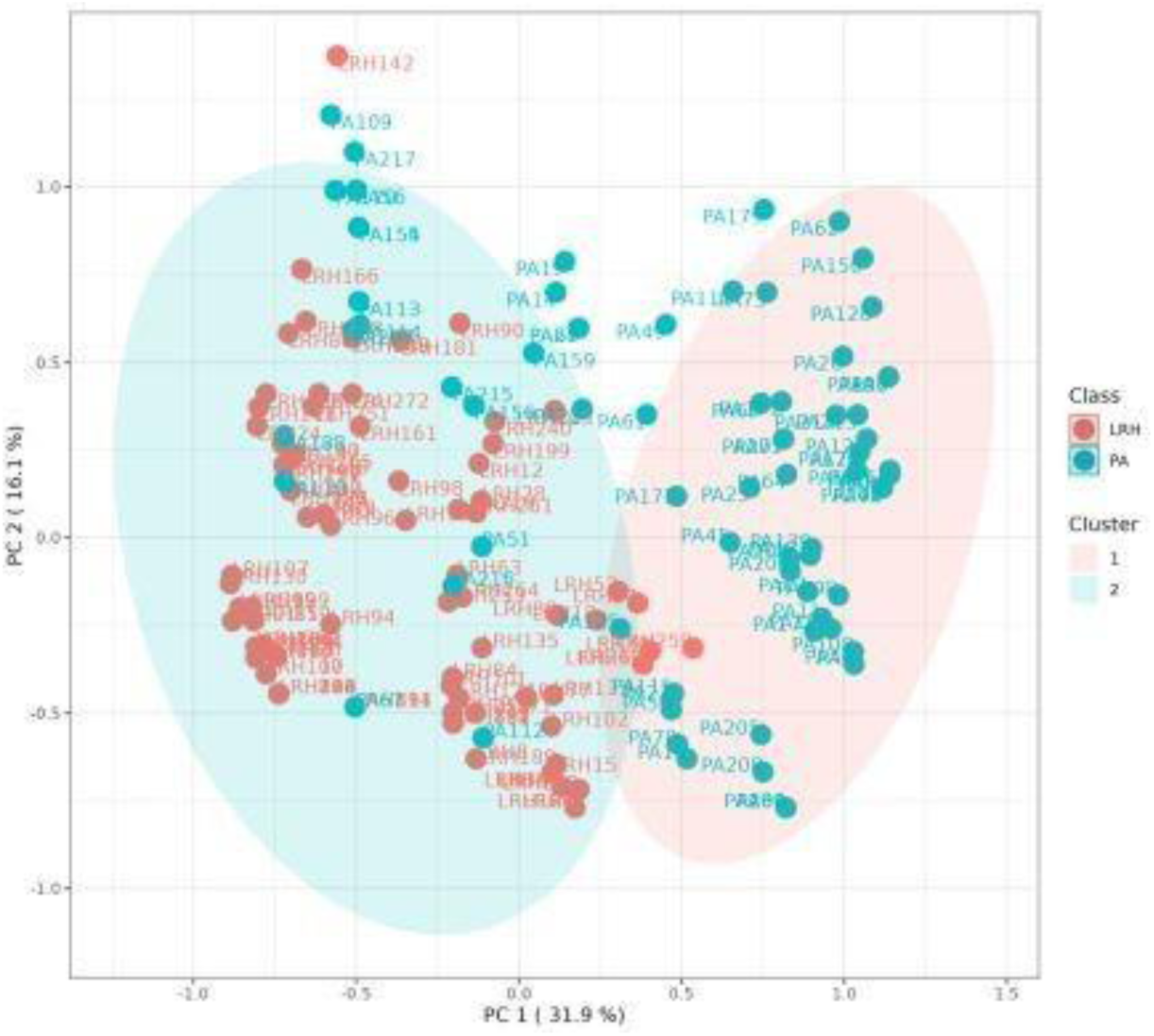
*k*-means cluster analysis (k=2) of the whole cohort. Each patient is represented by a dot with a corresponding ID label. Red dots: low-renin HTN (LRH) patients; green dots: primary aldosteronism (PA) patients). The 95% centroid confidence ellipses are shown for both clusters.

**Figure 4.**
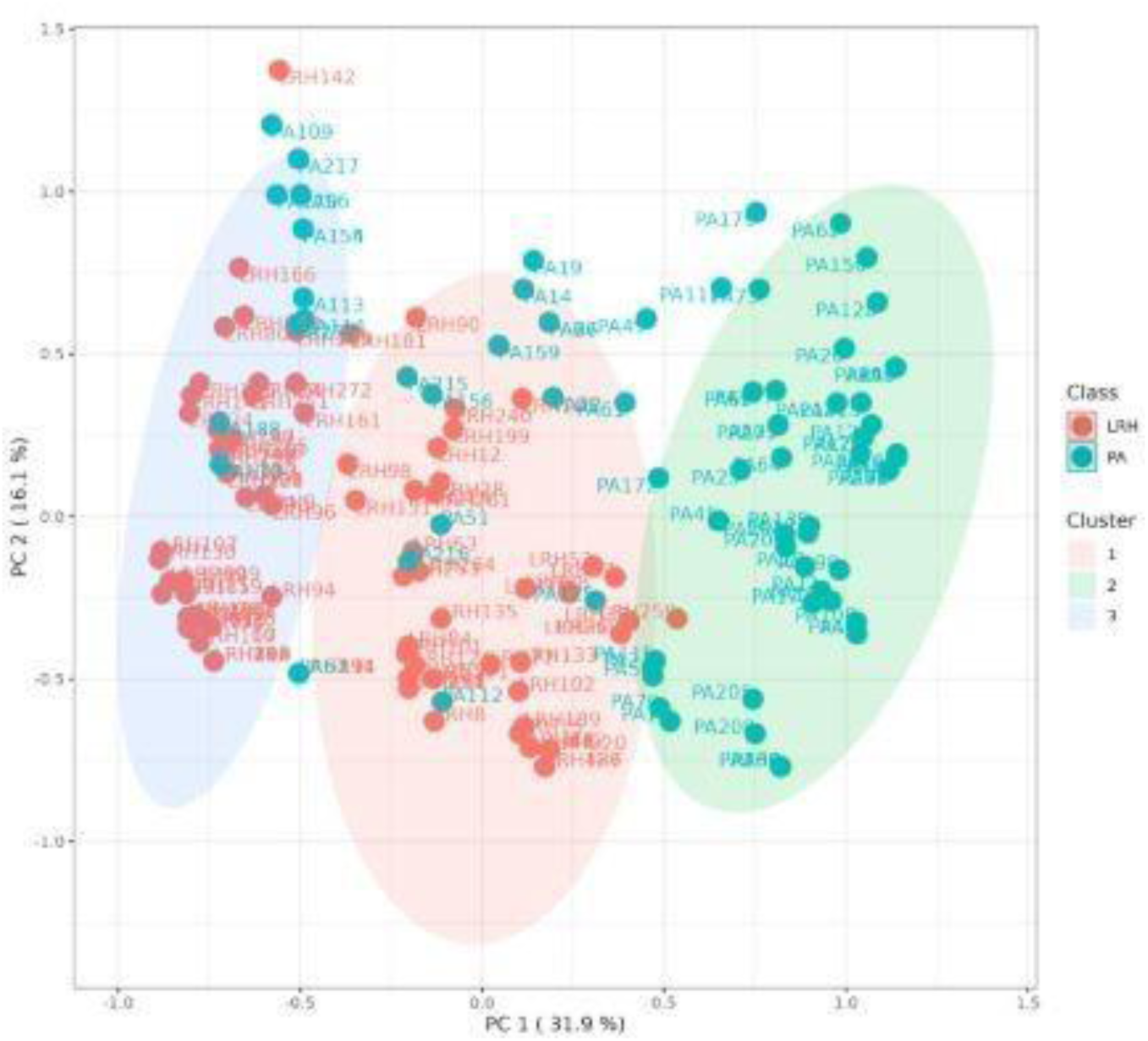
*k*-means cluster analysis (k=3) of the whole cohort. Each patient is represented by a dot with a corresponding ID label. Red dots: LRH (low-renin HTN) patients; green dots: primary aldosteronism (PA) patients. The 95% centroid confidence ellipses are shown for the *k*=3 clusters.

**Figure 5.**
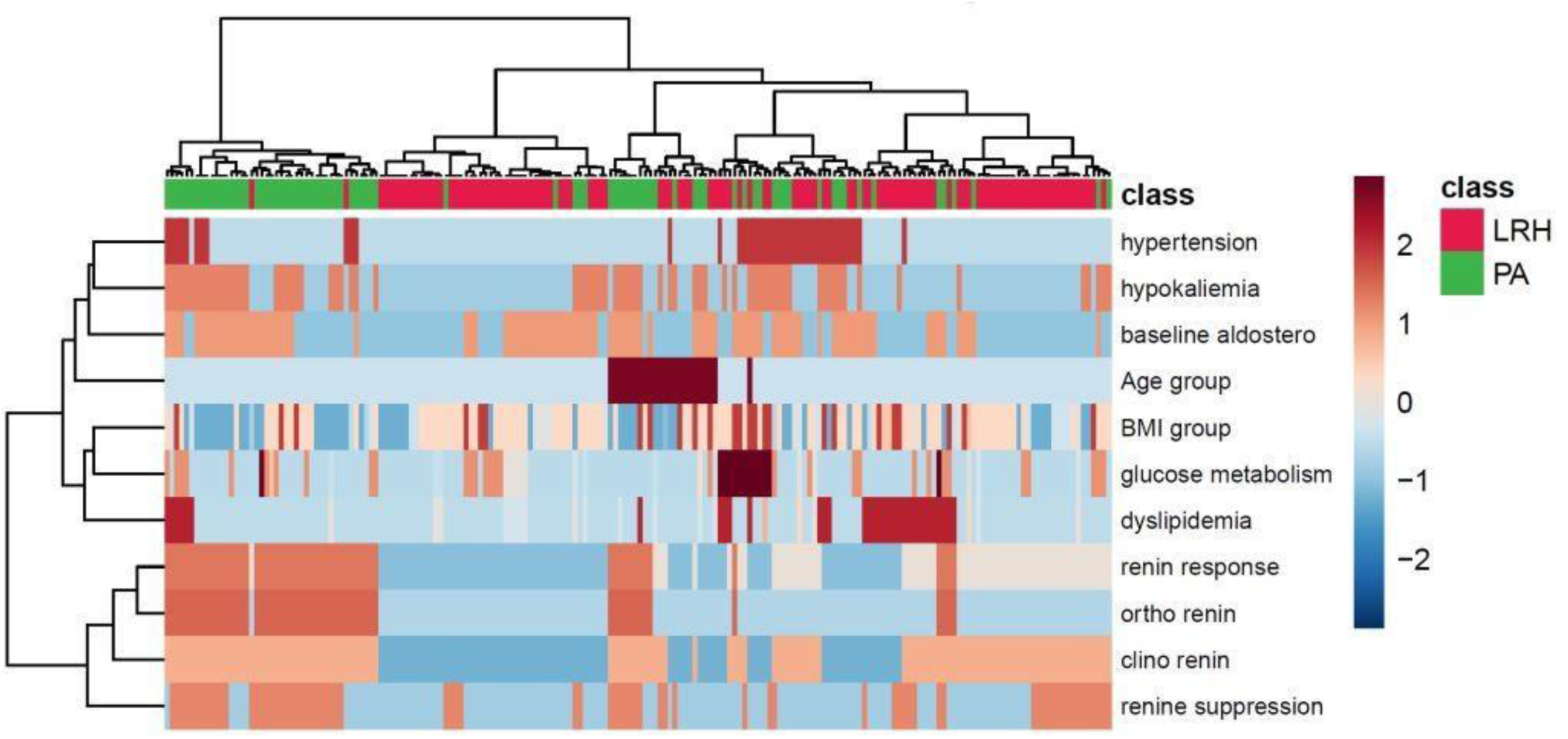
Hierarchical clustering shown as heatmap. Euclidean distance measure, Ward clustering method. Red dots: LRH (low-renin HTN) patients; green dots: primary aldosteronism (PA) patients.

**Table 4.** Clinical and biochemical picture of PA and low-renin HTN patients within Cluster 1 and Cluster 2. Data are presented as median and IQR (in brackets) or frequencies and percentages (in brackets). Abbreviations: BMI: body mass index, HTN: hypertension, GMA: glucose metabolism lterations. a) p < 0.05 vs Cluster 1; b) p < 0.001 vs Cluster 1; c) p < 0.01 vs Cluster 1

|  | PA (n=80) |  | Low renin HTN (n=110) |  |
| --- | --- | --- | --- | --- |
|  | Cluster 1<br>(n=57) | Cluster 2<br>(n= 23) | Cluster 1<br>(n=6) | Cluster 2<br>(n=104) |
| **Age** (years) | 51 (46-60.5) | 50 (46-62) | 57 (49-68) | 53 (46-63.2) |
| **Sex** (F/tot) | 25/57 (44%) | 12/23 (52%) | 6/6 | 64/104 (61%) |
| **Aldosterone baseline**<br>(pmol/L) | 755 (521-1116.7) | 1030 (687-1570)<sup>a</sup> | 474 (416-492.5) | 500 (401-674.7) |
| **Renin baseline**<br>(suppressed/tot) | 42/57 (74%) | 5/23 (22%)<sup>b</sup> | 4/6 (67%) | 23/104 (22%)<sup>a</sup> |
| **K<sup>+</sup>** (mmol/L) | 3.2 (2.8-3.6) | 3 (2.8-3.5) | 3 (2.8-3.4) | 3.9 (3.6-4.2)<sup>c</sup> |
| **HTN drugs** (≥ 3 drugs/tot) | 11/57 (19%) | 11/23 (48%)<sup>a</sup> | 0/6 | 15/104 (14%) |
| **GMA** (normal/tot) | 38/54 (70%) | 15/19 (79%) | 4/6 (67%) | 69/91 (76%) |
| **BMI** (kg/cm<sup>2</sup>) | 25 (22.7-28) | 26 (24.5-28.5) | 24 (23-31.5) | 27.5 (24-30) |
| **Renin in CP**<br>(suppressed/tot) | 57/57 (100%) | 8/23 (35%)<sup>b</sup> | 6/6 (100%) | 41/104 (39%)<sup>c</sup> |
| **Renin in OP**<br>(suppressed/tot) | 54/57 (95%) | 0/23<sup>b</sup> | 2/6 (33%) | 0/104<sup>a</sup> |
| **Renin response**<br>(always suppressed /tot) | 54/57 (95%) | 0/23<sup>b</sup> | 2/6 (33%) | 0/104<sup>b</sup> |
| **Aldosterone in CP**<br>(pmol/L) | 538 (376.5-804) | 573 (488.5-1037) | 274 (177-315) | 248.5 (179-373) |
| **Aldosterone in OP**<br>(pmol/L) | 860 (551.5-1218.5) | 1094 (698.5-1640) | 512.5 (406.5-676.2) | 628 (432.2-886.2) |

Abbreviations: BMI: body mass index, HTN: hypertension, GMA: glucose metabolism alterations. a) $p < 0.05$ vs Cluster 1; b) $p < 0.001$ vs Cluster 1; c) $p < 0.01$ vs Cluster 1

## Discussion

We studied the role of PST in the diagnostic work-up of patients with a high likelihood of PA, since the whole study population presented with arterial HTN and increased ARR. Nowadays, PA is considered a continuum spectrum of disease characterized by autonomous aldosterone secretion, ranging from mild to overt forms. Our cohort of low-renin HTN subjects showed normal-high aldosterone levels at diagnosis and therefore we may hypothesize that arterial HTN could be secondary to a mild aldosterone excess, which is not completely autonomous from renin levels and properly suppressed during PA confirmatory test. Given the increased cardiovascular morbidity and mortality of PA compared to essential HTN, it would be useful to rely on diagnostic tools with high accuracy in diagnosing overt forms of PA, which deserve a specific treatment (medical or surgical, according to the subtype) [25]. In the past PST was performed to distinguish unilateral to bilateral forms of PA. In a study by Fuss et al., patients with essential hypertension showed physiological response to PST with aldosterone increase mediated by angiotensin II. Bilateral PA showed the same behavior, while only 40% of unilateral PA showed aldosterone increase after PST. Therefore, the authors concluded that PST may improve PA subtyping in addition to AVS [14]. However, other studies showed conflicting results [26], [27], and PST was abandoned for this purpose due to low diagnostic accuracy and a significant overlap in responses. Nonetheless, in clinical practice, PST is easy and safe to perform; it does not require exogenous stimuli or dedicated staff and does not present peculiar contraindications as other confirmatory tests (i.e. SIT is contraindicated in case of severe uncontrolled HTN, renal insufficiency, heart failure, and CCT may lead to hypotension) [4]. Therefore, we aimed to study PST’s role in the diagnostic approach of patients with a high likelihood of PA, moving it from a test for PA subtyping (nowadays the indisputable gold standard is AVS) to a confirmatory test following a pathological ARR. Analyzing PST responses in our two groups (PA vs low-renin HTN patients) we noticed that the prevalence of suppressed renin levels in OP (after PST) was significantly higher in PA compared to low-renin HTN (69% vs 2%), while the majority of patients whose renin levels shifted from suppressed to measurable during PST or were always measurable belonged to the low-renin HTN group. These findings, and the demonstration that cortisol levels did not increase from CP to OP, highlight the physiological RAS response to postural stimulation in patients without PA. To corroborate this hypothesis, aldosterone levels during PST showed a higher rate of increase in low-renin HTN compared to PA (127% vs 50%), without differences in the two populations according to renin response from CP to OP. On the contrary overt PA, which is characterized by renin-independent autonomous aldosterone secretion, showed the persistence of suppressed renin levels in OP in most subjects. In a recent study [15] on the usefulness of PST in PA diagnosis, the authors reported that a DRCmax < 2 ng/L during PST was a predictor of PA. The authors also focused on aldosterone response during PST and found that 96% of PA showed a ≥50% increase in aldosterone levels during PST, while renin remained suppressed. This finding is partially in contrast to our results, since only 44% of our PA patients with always suppressed renin showed a ≥50% increase in aldosterone levels. We may speculate that it may be secondary to the different selection criteria: in the paper by Younes et al. controls were normotensive, and their number was 15% of the cohort. On the contrary, we considered a balanced group of PA and non-PA patients, and non-PA were not healthy controls but subjects with a high likelihood of being PA (hypertensive with pathological baseline ARR), therefore a group worthy to be tested for excessive aldosterone secretion. Regarding the aldosterone increase, they also concluded that there is no accurate aldosterone threshold during PST for predicting PA, in line with previous studies and also considering the known aldosterone intraindividual variability in PA. Concerning PST’s role in the diagnostic process of PA we agree that it should be placed in the setting of differential diagnosis of patients with suspected PA, as a confirmatory test. The non-hierarchical k-means clustering technique, when set to divide the entire cohort into two clusters, showed that most of the PA patients were correctly identified in Cluster 1 (57/80) while almost the entire cohort of low-renin HTN was identified in Cluster 2 (104/110). We then focused on the clinical picture of patients belonging to the two different clusters we observed a higher frequency of suppressed renin levels at baseline and during PST in Cluster 1 PA patients, and on the contrary lower potassium levels and a higher frequency of suppressed renin levels at baseline and during PST in Cluster 1 low-renin HTN patients. Therefore, an “hypokaliemic and renin suppressed phenotype” identified low-renin HTN patients of Cluster 1. Recent evidence suggested that overt PA is only the tip of the iceberg of the multidimensional spectrum of the condition, ranging from subclinical stages to severe forms, and from focal/multifocal to disseminated aldosterone producing areas affecting one or both the adrenal glands [25]. A renin-independent aldosterone excess was also found in mild hypertension and normokalemia, and even in normotensive subjects [6], [10]. In literature, the assessment of aldosterone levels in low-renin HTN showed a bimodal distribution and therefore, two pathological variants of this condition were proposed: an aldosterone-dependent form, which may be considered as a subgroup of PA, and a non-aldosterone-dependent form, which accounts for low renin HTN [28].Our findings, confirmed by the multivariate statistical analysis techniques, showed that PST had a good accuracy in the differential diagnosis of PA from low-renin HTN. In particular, renin response during PST and renin in OP were the more accurate predictive factors, combined with hypokalemia. Cluster analysis made it possible to better identify low-renin HTN compared to PA, confirming the continuum of the renin-independent aldosterone excess, though highlighting two subgroups within the entire cohort: a subclinical and an overt one. Renin response during PST allowed the identification of three distinct subgroups (always suppressed, from suppressed to measurable and always measurable) which were confirmed *a posteriori* by setting three clusters in the non-hierarchical k-means clustering technique, strengthening the data. Since the well-known presence of interfering conditions and the significant intra-individual variability in aldosterone, renin and ARR measurements, at least two ARRs are needed to exclude PA or proceed to further investigations [29]. Thus, in our opinion, the second ARR can be performed during PST: we support it to differentiate the two pathological conditions (PA vs low-renin HTN) ab initio, since it is an easy and safe to perform test. We are aware of the possibility of misclassifying cases based on the reference standard of confirmatory tests, in the absence of a gold standard for PA confirmation and considering the reduced performance rate of confirmatory tests especially in the “grey zone” of the continuous distribution of aldosterone excess from mild to overt forms. However, almost all patients of our cohort underwent two confirmatory tests to increase diagnostic accuracy and reduce misclassification. In particular, we decided to consider a strict cut-off to define positive SIT (aldosterone post-SIT> 10 ng/dl) and in case of aldosterone post-SIT in the “gray zone”, CCT response was evaluated to confirm or rule out PA. Based on this criterion, as shown in Figure 1, confirmatory tests clearly defined the “extremes” of disease distribution: overt PA with always suppressed renin levels during PST and positive response to confirmatory test (54/56) and low-renin HTN patients with always measurable renin levels and negative response to confirmatory tests (63/78). Unfortunately, regarding our cohort of low-renin HTN patients, follow up data are not available for all the 110 subjects since we are a tertiary referral center and thus, after the first evaluation for confirming the diagnosis, patients may be remitted to primary care setting, with the aim to refer to our center in case of worsening of the clinical profile. Nevertheless, we would like to underline that, in our opinion, interpreting PST response should not result in a dichotomous definition of PA vs non-PA, but rather an early identification of overt PA cases who deserve subtyping diagnosis (i.e. AVS). On the other hand, we would like to recommend that patients classified as low-renin HTN must receive an adequate follow up despite negative confirmatory tests, since they had at least one positive ARR and therefore they could evolve to overt PA in the following years. However, given negative confirmatory tests they don’t need to be straight referred to AVS, yet a medical treatment with MRAs must be considered. For this purpose, MRAs showed efficacy in blood pressure lowering in patients with low-renin HTN compared to other commonly used antihypertensive classes [30]. Another important point could be PST’s role in the follow-up of patients with a high likelihood to evolve to overt PA. A recent study by Buffolo *et al*. [30] on the long-term follow-up of patients with elevated ARR and negative confirmatory tests showed that about 20% of patients with a negative confirmatory tests develop overt PA after five years. Therefore, considering that aldosterone excess is a continuous condition, PST may be useful in the follow-up of patients with worsening of blood pressure control and/or decreasing of potassium levels, to early detect an incipient PA.

## Limitations

The main limitation of the study is its retrospective and monocentric design, which led to exclusion of some cases for incomplete data. Second, we did not prospectively compare the PST to a gold-standard confirmatory test for PA, because a gold standard with 100% diagnostic accuracy does not exist: a solid diagnosis of PA can be confirmed after adrenalectomy and the detection with immunohistochemistry of an aldosterone-producing (CYP11B2-positive) adenoma.

## Conclusions

To conclude, in a setting of hypertensive patients with at least one positive ARR and therefore suspected PA, PST showed high diagnostic accuracy in predicting PA vs low-renin HTN and this was confirmed by different analytical techniques, from the simplest to the elaborated ones. PLS-DA and PCA confirmed that renin in OP, renin response to PST, and presence of hypokalemia were the most relevant parameters to distinguish PA from low-renin HTN.

## Disclosure Summary / Declarations

### Authors’ contributions

Elena Pagin, Alberto Madinelli and Giorgia Antonelli: data collection; Irene Tizianel, Simona Censi and Eugenio Ragazzi: data analysis; Irene Tizianel and Filippo Ceccato: writing of original draft; Franco Mantero: supervision; Chiara Sabbadin and Mattia Barbot: writing-review and editing; Caterina Mian: funding acquisition. All authors meeting authorship criteria arelisted, and all authors certify that sufficient participation was given in order to take publicresponsibility for the content contained within this manuscript. All authors approved the final version of the paper. Research involving human participants and patient consent: Informed consent was obtained in all patients.

### Data availability statement

All data generated or analyzed during/ in this study are included in this published article, or in the repository data available online in the website of Padova University https://researchdata.cab.unipd.it/id/eprint/1508

